# Multiple Partially Overlapping Neural Modules Orchestrate Conflict Processing

**DOI:** 10.1101/2025.02.21.639534

**Authors:** Melinda Sabo, Manuel Varlet, Edmund Wascher, Patrick D. Gajewski, Tijl Grootswagers

**Affiliations:** Leibniz Research Centre for Working Environment and Human Factors, Dortmund, Germany; Yale University, Department of Psychology; The MARCS Institute for Brain, Behaviour and Development, Western Sydney University, Sydney Australia; School of Psychology, Western Sydney University, Australia; School of Computer, Data and Mathematical Sciences, Western Sydney University, Australia

**Author notes:** Correspondence concerning this article should be addressed to Melinda Sabo, Yale University, Department of Psychology, 100 College St. 06510 New Haven CT, USA.

**Keywords:** cognitive control, attention, Simon task, Stroop task, decoding, EEG

## Abstract

Cognitive conflict is a ubiquitous aspect of our daily life, yet its underlying neural mechanisms remain debated. Competing theories propose that conflict processing is governed by either a domain-general system, multiple conflict-specific modules, or an architecture involving partially overlapping mechanisms. The aim of the current study was to clarify how these mechanisms operate at the neural level using a data-driven decoding approach. We analyzed electroencephalogram (EEG) data from 507 healthy participants (ages 20–70) from the Dortmund Vital Study (ClinicalTrials.gov NCT05155397) who completed three conflict tasks: a Change Detection task, a Simon task, and a Stroop task. Using multivariate decoding to distinguish conflict from non-conflict trials, we observed robust within-task decoding across all tasks. However, cross-task decoding revealed shared conflict representations only in specific task pairs. These findings support an account, in which conflict processing relies on partially overlapping neural mechanisms rather than being fully domain-general or entirely task specific. We argue that conflict-related neural processes might be best conceptualized as a continuum of overlap and differentiation, with domain-generality and domain-specificity representing the endpoints of this spectrum.

**Research Transparency Statement:** The authors declare no conflicts of interest. Funding: This work was supported by the International Visiting Scholar Program at The MARCS Institute for Brain, Behaviour, and Development of Western Sydney University (MS), and the Australian Research Council grants DE230100380 (TG) and DP220103047 (MV).

<u>Artificial intelligence</u>: Artificial intelligence assisted technologies were utilized solely for language refinement and grammar corrections, ensuring clarity and readability without altering the original content or ideas. <u>Ethics</u>: The study was conducted in accordance with the Declaration of Helsinki and approved by the Institutional Ethics Committee of the Leibniz Research Centre for Working Environment and Human Factors, Ardeystraße 67 44139 Dortmund, Germany (IfADo; code A93-3) as detailed in the published study protocol (Gajewski et al., 2022). <u>Preregistration</u>: The current study was not preregistered. However, the dataset used in this manuscript originates from the Dortmund Vital Study, which is registered on ClinicalTrials.gov (NCT05155397). <u>Data</u>: A sample dataset is publicly available at https://doi.org/10.17605/OSF.IO/EDTNM <u>Analysis Scripts</u>: All analysis scripts are publicly available at https://doi.org/10.17605/OSF.IO/EDTNM.

## 1. Introduction

In our everyday life, we are constantly exposed to situations where we must overcome conflict. It is often the case that our cognitive system triggers an automatic response, while the situation requires a different, competing response. For example, if someone accustomed to driving in a country where vehicles travel on the left side of the road visits a country where driving occurs on the right side, they might instinctively look in the wrong direction when crossing the street. Overcoming this habitual response requires conscious effort and adjustment. This scenario exemplifies the type of cognitive conflict we must navigate. Such adaptations are made possible through cognitive control, a fundamental mechanism that allows humans to flexibly adjust to the ever-changing demands of the environment.

Conflict processing has been extensively studied in laboratory settings using various experimental paradigms. One prominent example is the Stroop task (Heidlmayr et al., 2020; MacLeod, 1991; Parris et al., 2022; Stroop, 1935), where participants are shown a color word (e.g., “red”) written in incongruent ink (e.g., blue). Because the color and word meaning are processed simultaneously, the irrelevant word meaning interferes with the display color and leads to conflict. Over time, researchers have offered different interpretations for the level at which conflict arises in the Stroop task (for a review, see MacLeod, 1991). Early perceptual-encoding accounts proposed that competition occurs during initial perceptual processing. However, the more widely accepted view holds that conflict emerges later, during response selection (for a review, see MacLeod, 1991). This is due to overlapping stimulus-response mappings. The prepotent but irrelevant stimulus-response set has to be inhibited, whereas the relevant one has to be activated, which delays the response execution in conflict trials (Gajewski et al., 2020). Another widely used paradigm is the Simon task, in which participants respond with the left or right hand based on a symbol’s identity. Conflict occurs when the symbol’s spatial location (e.g., appearing on the left) conflicts with the required response (e.g., right hand), creating response-level interference (Cespón et al., 2020; Hommel, 2011; Leuthold, 2011; Simon, 1969). Finally, conflict has also been examined in Change Detection paradigms, in which participants report changes in a specific feature across two successive stimulus displays (Schneider & Wascher, 2013; Wascher & Beste, 2010). Perceptual conflict arises when task-irrelevant changes compete with the relevant feature change, leading to reduced behavioral performance. Although conflict processing has been less frequently studied in these tasks compared to the Stroop or Simon paradigms, available evidence indicates similar behavioral patterns: accuracy and response time differences between conflict and non-conflict trials are reliably observed (Schneider et al., 2012a, 2012b, 2014; Schneider & Wascher, 2013; Wascher & Beste, 2010). Moreover, these studies report established EEG markers of conflict resolution, including fronto-central N2 modulations and the N2pc component linked to the selection of task-relevant information (Schneider et al., 2012a, 2012b, 2014; Schneider & Wascher, 2013; Wascher & Beste, 2010).

The neural mechanisms underlying conflict processing and control remain the focus of ongoing debate. One influential account proposes that conflict is processed in a domain-general manner, such that different types of conflict engage a common neural system. Neuroimaging studies repeatedly report overlapping activation in regions such as the supplementary motor area, the anterior cingulate cortex, and the anterior insula (Chen et al., 2018; Vermeylen et al., 2020). In particular, the ACC and anterior insula have been linked to detecting conflict and signaling the need for cognitive adjustment across both cognitive and emotional tasks (Chen et al., 2018). Behavioral evidence also supports this view through the congruency sequence effect, the reduction of conflict effects (e.g., faster responses, improved accuracy) following previous conflict trials. When this effect transfers across tasks or conflict types, it is often interpreted as evidence for domain-general conflict processing and control. Numerous studies have reported such transfer, showing that congruency sequence effects can generalize across a wide range of tasks and conflict domains (Braem et al., 2014; Fernandez-Duque & Knight, 2008; Freitas et al., 2007; Freitas & Clark, 2015; Funes et al., 2010; Kan et al., 2013; Kleiman et al., 2014; Schmidt & Weissman, 2014).

While domain-general models propose a unified neural mechanism for conflict detection and control, accumulating evidence has called this assumption into question. For example, Kan and colleagues (2013) reported that the congruency sequence effect could transfer from sentence-reading and perceptual detection tasks to the Stroop task. However, two subsequent studies failed to replicate this finding (Aczel et al., 2021; Dudschig, 2022), casting doubt on the generalizability of the effect. A broader set of studies further challenged this view, showing that congruency sequence effects are often limited to the same task or conflict type, with little evidence for transfer across distinct tasks (Akçay & Hazeltine, 2011; Boy et al., 2010; Braem et al., 2014; Egner et al., 2007; Forster & Cho, 2014; Funes et al., 2010; Kim et al., 2012; Kunde et al., 2012; Kunde & Stöcker, 2002; Rünger et al., 2010; Schlaghecken et al., 2011; Verbruggen et al., 2005; Wendt et al., 2006; Wühr et al., 2015; Zhu et al., 2025). Other researchers have adopted a more nuanced view, acknowledging partial transferability: observed in some contexts but not universally across tasks or conflict types (Freitas & Clark, 2015).

Neuroimaging studies have mirrored these behavioral findings, indicating that domain-general accounts may not fully capture how conflict is processed (e.g., Li et al., 2017). For instance, Jiang and Egner (2014) used fMRI to compare stimulus and response conflict and found that domain-specific activation patterns were more robust than domain-general ones, interpreting this as evidence for a hybrid architecture. Likewise, Fu and colleagues (2022) reported that the medial frontal cortex encodes both shared and conflict-specific signals. Additional work further supports this differentiation: stimulus-based conflicts tend to recruit frontal and parietal regions, whereas response-based conflicts primarily engage motor and premotor areas (Cespón et al., 2020; Egner, 2008; H. Li et al., 2019; Parris et al., 2019, 2022; Zmigrod et al., 2016). Together, this growing body of research challenges the notion of a fully domain-general conflict resolution system and suggests that domain-specific mechanisms (see Egner, 2008 for a review) also play a substantial role.

Building on this prior work, the current study aims to clarify whether conflict processing is best described as (i) a domain-general neural mechanism, (ii) a set of conflict-specific modules, or (iii) partially overlapping neural mechanisms, which entail neural dynamics that are neither entirely shared nor fully task-specific. Although related questions have been examined previously, this study makes three novel contributions to the literature. First, we use a large, representative sample, addressing a major limitation of prior studies, where interpretations of null effects (e.g., failures of the congruency sequence effect to transfer across tasks; Aczel et al., 2021; Dudschig, 2022), have often been constrained by small sample sizes and potentially limited statistical power. Second, we adopt a data-driven multivariate approach capable of detecting spatiotemporal regularities in the EEG signal without requiring strong a priori assumptions. Third, by using EEG, we capture the rapid temporal dynamics of conflict processing, which remain difficult to assess with fMRI-based approaches that have dominated this area (e.g., Jiang & Egner, 2014; Vermeylen et al., 2020), leaving the temporal dynamics of conflict processing comparatively underexplored (for an exception, see Fu et al., 2022). Together, these methodological strengths enable a more robust and temporally precise test of whether conflict processing is domain-general, conflict-specific, or it relies on partially overlapping neural mechanisms.

In the current study, EEG and behavioral data were collected from 507 participants, spanning an age range of 20–70 years. Data were part of the Dortmund Vital Study (ClinicalTrials.gov NCT05155397; see study protocol Gajewski et al., 2022). Participants performed three cognitive tasks involving conflict processing. A Change Detection task entailing task-relevant and irrelevant features, a Simon task, and a Stroop task (see Figure 1). Using multivariate pattern analysis, we examine whether a linear classifier can learn to distinguish conflict from non-conflict trials based on EEG data. First, we train a classifier to identify conflict-related patterns within each task. We not only run this analysis in the time domain, but also in the frequency domain, as previous research highlights the role of theta-band activity in conflict processing (Hanslmayr et al., 2008; Nigbur et al., 2011). Next, we investigate whether these conflict signals generalize across tasks by training the classifier on one task and evaluating its performance on the remaining task, a process referred to as cross-task decoding. In this context, three outcomes are possible. The *domain-general conflict processing perspective* predicts above-chance decoding performance for both within-and cross-task decoding across all tasks. Alternatively, under the *multiple conflict-specific modules* framework, only within-task decoding is expected to produce statistically reliable results.

**Figure 1.**
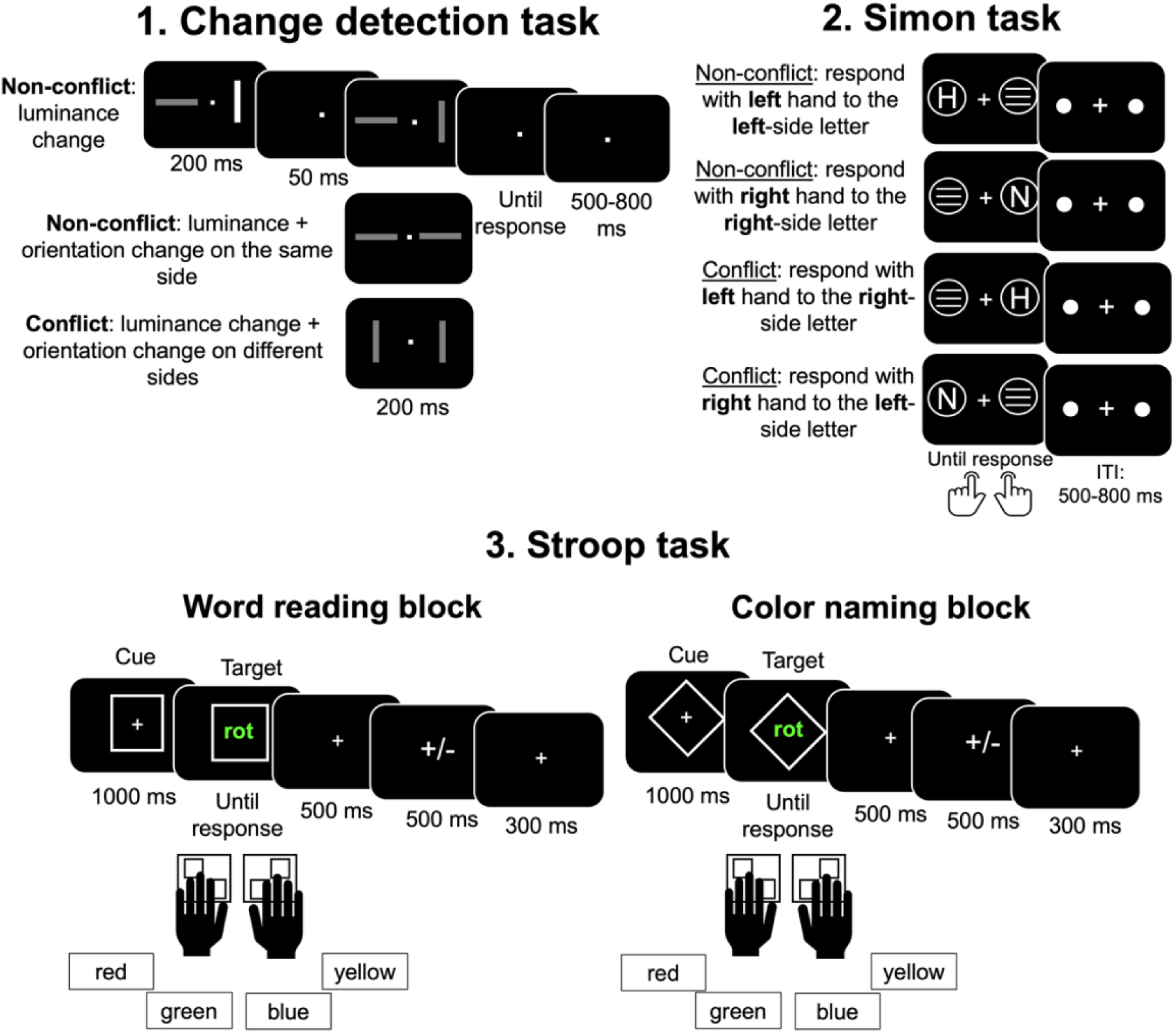
Overview of experimental tasks involving conflict at different levels. **(a) Change Detection task**: Participants viewed two squares on either side of a fixation cross. Stimuli could include a luminance change (non-conflict), a luminance and orientation change on the same side (non-conflict), or a luminance and orientation change on opposite sides (conflict). Participants responded to the change location, with trials separated by an inter-trial interval (ITI) of 500–800 ms. **(b) Simon task**: Participants responded to a letter presented on either side of the screen (H or N) using a spatially compatible or incompatible hand. Non-conflict trials required spatially congruent responses, while conflict trials involved spatially incongruent responses, with an ITI of 500–800 ms. **(c) Stroop task**: Two blocks of trials were included: a **Word Reading Block**, in which participants read the word regardless of its ink color and a **Color Naming Block**, in which participants named the ink color of a word regardless of the word’s meaning. Color Reading and Naming blocks were not mixed and were completed as separate blocks. The Word Reading Block served as a baseline level, as conflict in these trials was reduced. The task included four colors, red, yellow, green, blue, displayed in German (i.e., “rot”, “gelb”, “grün”, “blau”). Responses were made using four keys corresponding to the four possible colors, and each trial began with a cue indicating the block type (reading or naming), followed by the target word and a variable ITI.

Finally, the *partially overlapping neural mechanisms account* entails an above-chance decoding accuracy for the within-task procedure, and for some, but not all cross-task decoding combinations. To foreshadow our findings, results showed that conflict decoding was successful within individual tasks. Cross-task decoding analyses indicated that the conflict signal only partially generalized across tasks, supporting the predictions of the *partially overlapping neural mechanisms account*.

## 2. Results

### 2.1. Behavioral results

First, as a sanity check, we verified that the classic conflict effects reported in previous studies also appear in our sample. Figure 2 shows the accuracy and response time distributions for conflict versus non-conflict trials across all tasks. Both accuracy and response times differed reliably between conflict and non-conflict conditions, with Bayes factors suggesting very strong evidence in favor of the effect (Figure 2, Table 1 and 2). The only exception was accuracy in the Stroop Reading task, which did not show a significant conflict vs. non-conflict difference.

**Figure 2.**
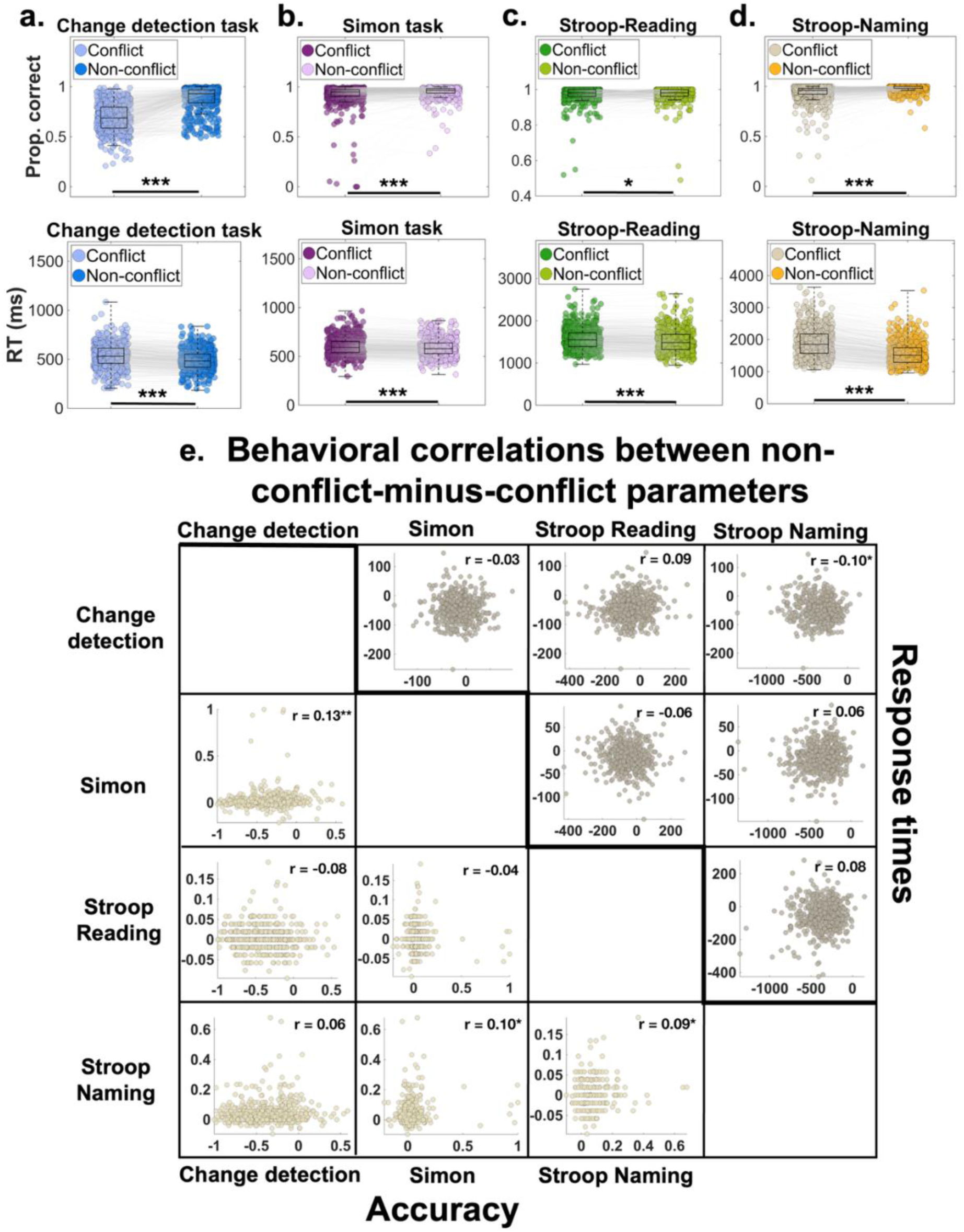
Behavioral performance and cross-task correlations. **a–d**. Accuracy (top panels) and response times (bottom panels) for the four tasks: **a.** Change Detection task, **b.** Simon task, **c.** Stroop Reading task, and **d.** Stroop Naming task. In each panel, boxplots summarize the distribution of performance for conflict versus non-conflict trials. The central line of each boxplot indicates the median; the lower and upper edges represent the 25th and 75th percentiles, respectively. Whiskers denote the most extreme non-outlier values, and individual participant means are plotted as points. Gray lines connect data points from the same participant across conditions. “Prop. correct” stands for proportion correct, which is the accuracy measure obtained by subtracting conflict from non-conflict trials. More positive values indicate higher performance. “RT” stands for response times. **e.** Pairwise correlations between non-conflict-minus-conflict difference scores across tasks for accuracy (lower triangle) and response times (upper triangle). Each scatterplot displays one participant per point, with corresponding Spearman correlation coefficients shown in each panel (\**p* < 0.05; \*\**p* < 0.01; \*\*\**p* < 0.001).

**Table 1:** Descriptive statistics.

Mean and standard deviation (SD)
| Task | Condition | N | Mean accuracy | SD accuracy | Mean response times | SD response times |
| --- | --- | --- | --- | --- | --- | --- |
| Change Detection | conflict | 507 | 0.68 | 0.14 | 534.40 ms | 120.56 ms |
| Change Detection | non-conflict | 507 | 0.87 | 0.12 | 488.50 ms | 106.89 ms |
| Simon | conflict | 507 | 0.91 | 0.12 | 602.86 ms | 97.18 ms |
| Simon | non-conflict | 507 | 0.95 | 0.06 | 588.76 ms | 95.34 ms |
| Stroop Reading | conflict | 507 | 0.97 | 0.04 | 1572.35 ms | 263.76 ms |
| Stroop Reading | non-conflict | 507 | 0.97 | 0.03 | 1518.18 ms | 275.12 ms |
| Stroop Naming | conflict | 507 | 0.93 | 0.09 | 1915.23 ms | 461.48 ms |

**Mean and standard deviation (SD)**
| Task | Condition | N | Mean accuracy | SD accuracy | Mean response times | SD response times |
| --- | --- | --- | --- | --- | --- | --- |
| Stroop Naming | non-conflict | 507 | 0.97 | 0.03 | 1564.18 ms | 370.20 ms |

**Table 2.** Conflict vs. non-conflict comparisons across tasks.

of the frequentist and Bayesian statistics for conflict vs. non-conflict comparisons
| Task | Behavioral parameter | df | *t* | *p* | *d<sub>av</sub>* | 95% CI | *BF<sub>10</sub>* |
| --- | --- | --- | --- | --- | --- | --- | --- |
| Change Detection | accuracy | 506 | -29.13 | < .001 | 1.40 | [-0.20, -0.17] | $3.13 \times 10^{106}$ |
| Change Detection | response times | 506 | 24.06 | < .001 | 0.40 | [42.15, 49.65] | $1.47 \times 10^{82}$ |
| Simon | accuracy | 506 | -6.14 | < .001 | 0.34 | [-0.04, -0.02] | $3.71 \times 10^6$ |
| Simon | response times | 506 | 10.85 | < .001 | 0.14 | [11.47, 16.54] | $3.82 \times 10^{21}$ |
| Stroop Reading | accuracy | 506 | -2.43 | 0.01 | 0.07 | [-0.005, -0.0006] | 0.93 |
| Stroop Reading | response times | 506 | 13.80 | < .001 | 0.20 | [46.45, 61.87] | $4.31 \times 10^{33}$ |
| Stroop Naming | accuracy | 506 | -14.00 | < .001 | 0.73 | [-0.05, -0.04] | $3.39 \times 10^{34}$ |
| Stroop Naming | response times | 506 | 42.35 | < .001 | 0.84 | [334.77, 367.33] | $3.20 \times 10^{164}$ |

Second, we computed the non-conflict-minus-conflict difference scores for accuracy and response times and examined their correlations across tasks. As shown in Figure 2e, four correlations reached significance: (i) an accuracy correlation between the Simon task and Change Detection task, *r*(506) = 0.13, *p* = .003; (ii) an accuracy effect between the Simon task and Stroop Naming task, *r*(506) = 0.10, *p* = .02; (iii) an accuracy effect between the Stroop Naming task and Stroop Reading task, *r*(506) = 0.09, *p* = .04; (iv) a response time correlation between the Change Detection task and Stroop Naming task, *r*(506) = –0.10, *p* = .02. All other correlations were small and non-significant (Figure 2). Together, these results indicate that conflict effects generalize only weakly across tasks.

### 2.2. Successful within-task decoding – evidence from the time domain

The first step in our decoding analysis was to identify conflict-related signals within each task. To achieve this, we trained a linear classifier to distinguish between conflict and non-conflict trials separately for each task. This analysis was performed at each time point across the trial. As shown in Figure 3, the classifier successfully differentiated between conflict and non-conflict trials. Statistical analyses revealed the following significant clusters: (i) 96–1396 ms for the Change Detection task; (ii) smaller clusters ranging 124-188 ms, and a sustained cluster between 224–1372 ms for the Simon task; (iii) smaller clusters between 720-1516 ms and a sustained cluster ranging 1524–1996 ms for the Stroop Reading task; (iv) 480–1112 ms for the Stroop Naming task. The right panels of Figure 3 show that Bayes factors provide very strong support for the effects at the significant time points. Figure 3 also highlights the range in which 95% of the response times for conflict and non-conflict trials occurred. Although there is some overlap between these time windows and the significant clusters, it is unlikely that response time differences between conflict and non-conflict trials are the primary driver of these decoding effects. This assertion is supported by the topographical maps for these critical time windows, which highlight a predominantly centro-frontal localization of the effect. Notably, these maps closely resemble those reported in the existing literature (Gajewski & Falkenstein, 2015; Heidlmayr et al., 2020; Schneider et al., 2012a).

**Figure 3.**
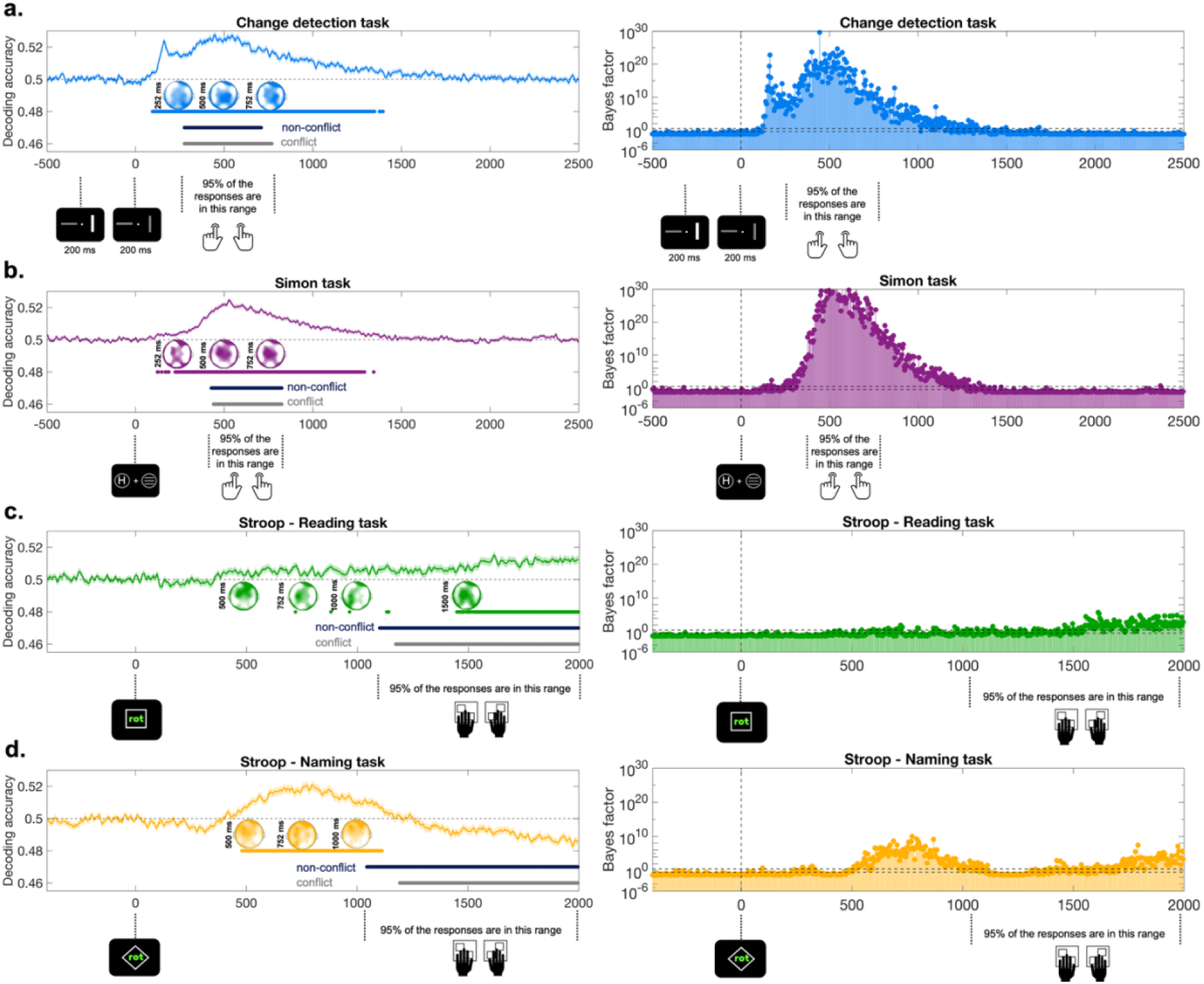
Decoding accuracy across time within each task a. Change Detection task, b. Simon task, and c, d. Stroop Reading and Naming tasks. The panels on the left side depict the decoding accuracy across time, while the ones on the right side illustrate the Bayes factors across time. Dashed lines at 0.5 on the left side panels indicate chance level, while the shaded area around the average decoding accuracy represents the standard error of the mean. Blue, purple, yellow, and green dotted lines denote significant clusters identified through statistical analyses. Black and gray dotted lines indicate the time ranges during which 95% of the response times for conflict and non-conflict conditions occur. Each time point is annotated with the task-specific events. The dotted lines at 3 and 0.33 on the right-side panels mark the area, where Bayes factors are inconsistent. All values above or below this indicate at least substantial evidence for the alternative or null hypothesis. The topographical maps within each task panel were derived from searchlight analyses conducted at the specific time points indicated next to each map. The decoding accuracy limits for the topographical maps are as follows: **a.** Change Detection task: 0.50–0.514; **b.** Simon task: 0.50–0.515; **c.** Stroop Reading task: 0.50–0.508 **d.** Stroop Naming task: 0.50–0.508.

### 2.3. Successful within-task decoding – evidence from the frequency domain

In a second step, we implemented an alternative within-task decoding analysis, which uses power values from different frequencies as a feature alongside electrodes. We ran a fast Fourier transform on the 0-1000 ms window following stimulus presentation and obtained power values for the frequency range between 1-50 Hz. We implemented this second within-task decoding procedure to determine whether power values from specific frequency ranges significantly contribute to decoding accuracy (Hanslmayr et al., 2008; Nigbur et al., 2011). Within each participant, the analysis resulted in a single decoding accuracy value.

To assess the statistical reliability of the results, we generated a null distribution of decoding accuracies by randomly shuffling conflict labels within each participant. The mean decoding accuracy was subsequently compared to the null distribution to determine its percentile rank. A value above the 95th percentile indicated significant above-chance decoding accuracy, while a value below this threshold was considered evidence for chance-level performance, supporting the null hypothesis. We obtained the following results: for the Simon task, Change Detection task, and Stroop Naming task, the mean decoding accuracy was at the 100th percentile of the null distribution, demonstrating robust evidence for above-chance decoding performance (Figure 4). In contrast, for the Stroop Reading task, the decoding accuracy was at the 67th percentile, providing evidence that the classifier could not reliably differentiate between conflict vs. non-conflict trials (Figure 4). These results were also supported by the Bayesian analysis, which revealed very strong support for the effect detected in the Change Detection task, Simon task, and Stroop Naming task. Nevertheless, the evidence was inconclusive for the Stroop Reading task (Figure 4).

**Figure 4.**
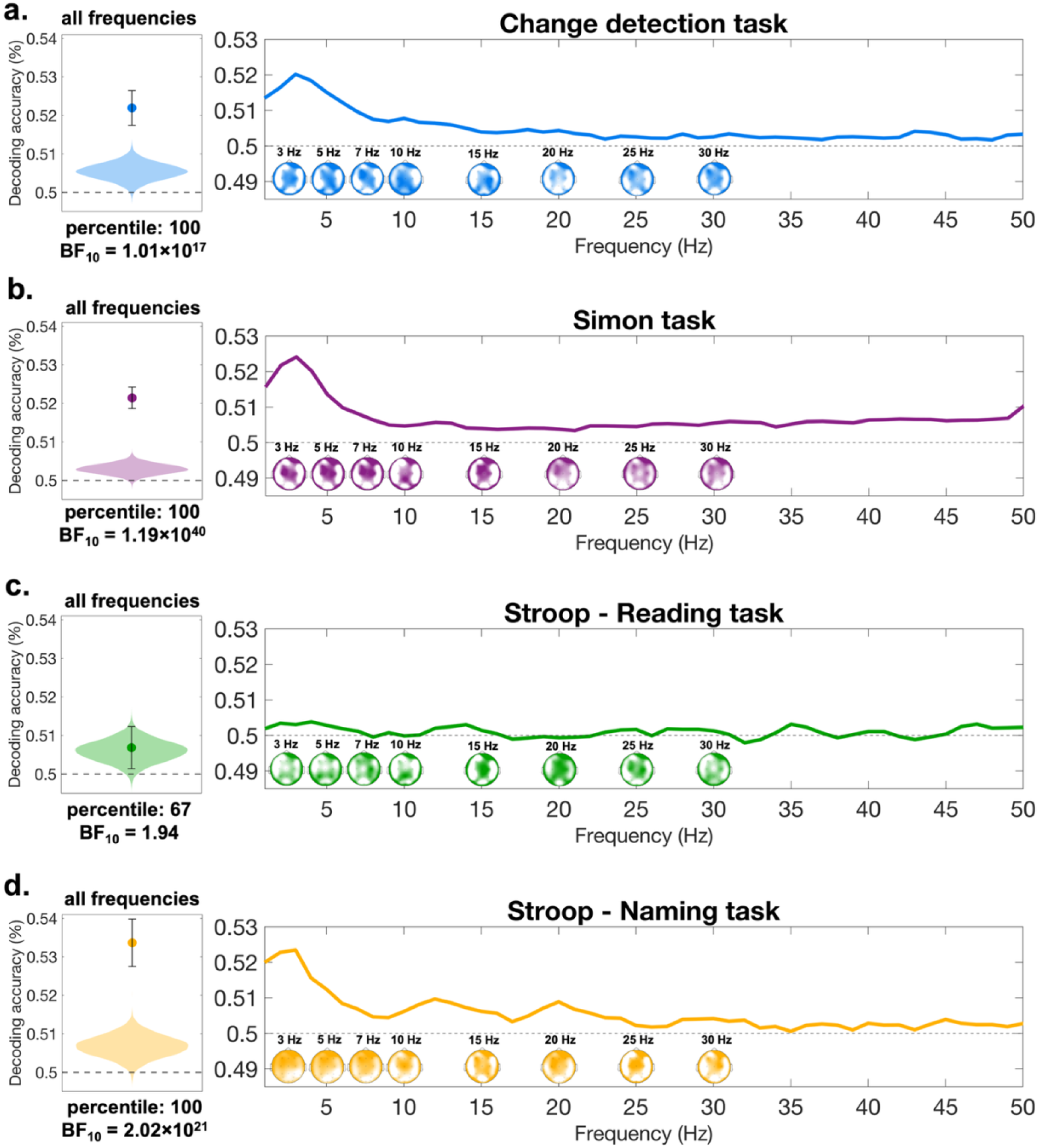
Within-task decoding based on frequency-domain information. Panels **a–d.** show the results of the second within-task decoding procedure performed on the power values between 1–50 Hz. The left-side panels display the comparison of mean decoding accuracy relative to the null distribution. The dashed gray line represents chance level, while the violin plot illustrates the null distribution generated by permuting conflict labels within each participant, with 10,000 permutations randomly sampled and averaged. As explained in Section 2.2, the null distribution values are not symmetrically centered around chance level. The dot represents the mean decoding accuracy, and the error bar indicates the confidence intervals of the mean. For panels **a., b., d.**, the average decoding accuracy corresponds to the 100th percentile, while in panel **c.**, it falls at the 67th percentile. Bayes factors are also reported. Comparisons **a., b., d.** are supported by very strong evidence favoring the alternative hypothesis, while the evidence for the Stroop Reading task remains inconclusive. The right-side panels **a–d.** depict the decoding accuracy from the searchlight analysis across frequency and electrode space. In the frequency representation, decoding accuracy values were averaged across all electrodes. The limits of the topographical maps are not uniform and vary across different maps. The minimum is at 0.50 and the maximum ranges between 0.503-0.51.

As shown in Figure 4 (left panels), the null distributions do not center around 0.50 but instead fall consistently above 0.50 across all four distributions. This pattern aligns with prior research suggesting that classification accuracies under the null hypothesis are not symmetrically distributed around chance (Allefeld et al., 2016).

Finally, we also conducted a searchlight analysis to examine whether specific frequency bands contributed more significantly to the decoding results. The analysis revealed that the highest decoding accuracy consistently occurred in the theta frequency range (3–7 Hz), which aligns with previous findings (Hanslmayr et al., 2008; Nigbur et al., 2011). Moreover, the topographical representation of decoding accuracies in the theta-band also revealed that these effects are maximal at fronto-central regions. This corresponds to the location where the conflict-related N2 reaches its maximum, previously localized to the anterior cingulate cortex (Ullsperger & Von Cramon, 2006; Vanveen & Carter, 2002; Yeung & Cohen, 2006).

Overall, the results from both procedures demonstrated that, within each task, conflict and non-conflict trials exhibit distinct EEG patterns, allowing for successful classification. The only exception was the Stroop Reading task. However, this is not surprising, as the reading task served as a baseline task and it did not involve conflict processing.

### 2.4. Partially generalizable conflict signal across tasks – evidence from time generalization analyses

The cross-task decoding analysis was conducted to determine whether neural patterns associated with conflict processing generalize across different cognitive tasks. Notably, the Stroop Reading task was excluded from this set of analyses due to its lack of robust evidence for within-task conflict decoding. We performed a time-resolved cross-task decoding, where we trained the classifier at each time point in one task and tested it at every time point. As shown in Figure 5, we observed a shared decodable conflict signal between the Change Detection task and Stroop Naming task. When the classifier was trained on the Change Detection task, a generalizable signal emerged at ∼248 ms, persisted until ∼1024 ms, and transferred to the ∼304-1992 ms window in the Stroop Naming task. A similar pattern appeared in the reversed analysis: training on Stroop Naming task yielded a generalizable signal beginning at ∼288 ms, lasting until ∼1992 ms, and transferring to the ∼312-904 ms interval in the Change Detection task.

**Figure 5.**
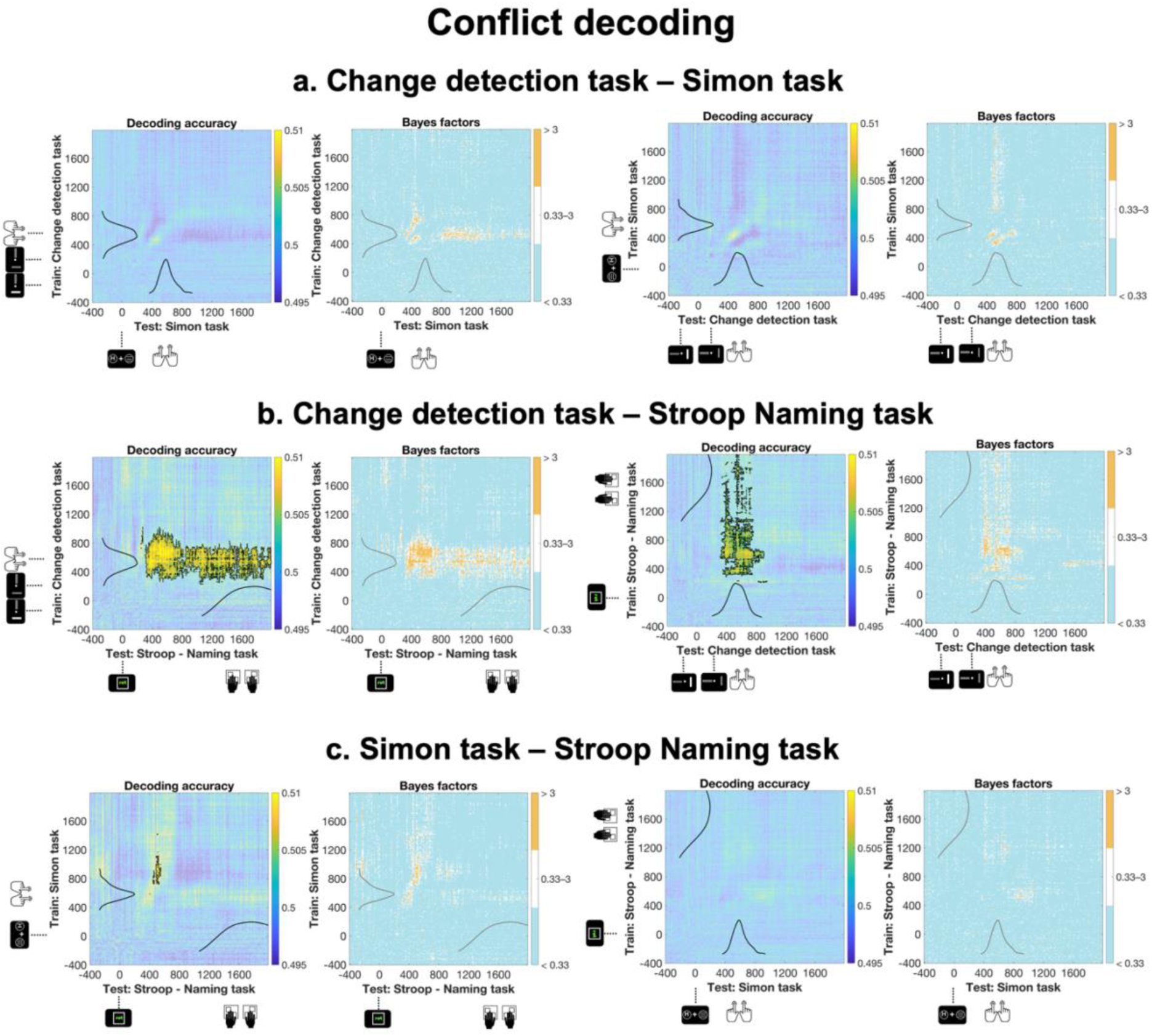
Temporal generalization of conflict decoding across tasks. Mean temporal generalization matrices showing cross-task decoding accuracy (left panels) and corresponding Bayes Factors (right panels) for all task pairings: **a.** Change Detection task and Simon task, **b.** Change Detection task and Stroop Naming task, and **c.** Simon task and Stroop Naming task. In all decoding accuracy plots, light blue indicates chance-level temporal generalization accuracy, whereas warmer colors (yellow to orange) reflect above-chance decoding performance. Significant clusters, identified via cluster-based permutation testing, are outlined by black contours in the decoding accuracy plots. In all Bayes Factor plots, orange indicates Bayes Factors above 3 (at least substantial evidence for the alternative hypothesis), white depicts inconclusive evidence (Bayes factors between 0.33-3) and cooler blue shades indicate Bayes Factors below 3 (at least substantial evidence for the null hypothesis). All panels also display the response time distributions which demonstrate that the temporal profile of the significant decoding clusters does not overlap directly with the response time distributions, indicating that the observed cross-task decoding is unlikely to be fully driven by motor-related activity.

We also identified a small significant cluster when training on the Simon task and testing on Stroop Naming task (∼720-1120 ms and ∼488-552 ms), though this effect did not replicate did not replicate when training and testing were reversed. No generalizable patterns were found between the Simon task and Change Detection task. All significant clusters were supported by at least substantial evidence, as indicated in the right panels of Figure 5. Importantly, to demonstrate that the lack of cross-task decoding in the Simon task and Change Detection task is not due to confounding factors, we ran a control analysis, in which we show that response hand (left vs. right) can be decoded with this technique across all task combinations (see Supplementary materials, Figure S1). Additionally, as indicated in Figure 5, which depicts the response time distributions within each task, significant decoding clusters do not overlap directly with responses. This suggests that the observed cross-task decoding is unlikely to be fully driven by motor-related activity.

In summary, these results suggest that conflict-related neural patterns generalize across some tasks but not others. Crucially, the absence of cross-task decoding in certain pairs cannot be attributed to methodological limitations, as certain features, such as response hand signals, generalized reliably across task combinations. Taken together, these findings support the partially overlapping neural mechanisms account.

## 3. Discussion

The primary aim of the current study was to investigate whether conflict processing is governed by (i) a domain-general neural mechanism; (ii) multiple conflict-specific modules; or (iii) it is best explained by partially overlapping neural mechanisms. To explore this, we examined whether a linear classifier could distinguish conflict from non-conflict trials based on EEG data across three distinct conflict tasks. Analyses were performed both within each task as well as across tasks. A key strength of the cross-task decoding approach lies in its ability to bypass assumptions about specific neural components or regions shared across tasks (Grootswagers et al., 2017; Hebart & Baker, 2018). Instead, the method leverages the classifier’s capacity to detect regularities in the data that might reflect task-independent conflict processing. By focusing on data-driven patterns, this study offers a robust approach to probing the generalizability of conflict signals, contrasting with prior research that often relies on predefined neural markers or brain regions of interest. Another distinctive aspect of this work is the dataset used for these analyses. Unlike most prior studies, which often rely on smaller or less diverse samples, this study drew on a large, representative sample of 507 individuals aged from 20 to 70 years. This large and representative sample enhances the generalizability of our findings, providing insights into conflict processing across a wide cross-section of the population. The size and diversity of this dataset mark a significant methodological advance for robust and reproducible investigations of conflict processing and control and its neural correlates.

Our findings revealed that the classifier reliably distinguished conflict from non-conflict trials across all tasks. These results align with prior research highlighting distinct neural signatures of conflict processing and control. Specifically, the fronto-central N2 component has been consistently linked to conflict monitoring in Simon task, Stroop task, and Change Detection task (Gajewski & Falkenstein, 2015; Heidlmayr et al., 2020; Wascher & Beste, 2010). The parietal P3 component is another EEG correlate, whose modulation has been frequently shown in conflict tasks (Ila & Polich, 1999; Leuthold, 2011; Polich, 2007). Additionally, theta-band oscillatory activity (∼3–7 Hz) has also been strongly associated with conflict resolution, particularly in Stroop and Simon tasks (Hanslmayr et al., 2008; Nigbur et al., 2011). Consistent with these findings, our searchlight analysis found that within-task decoding accuracy peaked at 3 Hz and extended across the theta range. Together, these results provide robust evidence that conflict processing can be neurally tracked within each task, with components such as the N2, P3, and theta oscillations possibly underlying the successful decoding.

The cross-task decoding results revealed partial generalizability of the conflict signal across tasks. We observed a robust shared signal between the Change Detection task and Stroop Naming task, and a weaker but still reliable effect between the Stroop Naming task and Simon task. In contrast, no cross-task decoding emerged between the Change Detection task and Simon task, a conclusion further supported by at least substantial Bayes factors. This pattern challenges a strictly domain-general account, which would predict cross-task decoding for all task pairs as long as each task contains a decodable conflict signal individually. At the same time, the findings are also difficult to reconcile with a fully domain-specific framework, which assumes that conflict processing relies on highly task-specific neural patterns and therefore should not produce any cross-task generalization. Instead, the results point toward an account entailing partially overlapping neural mechanisms.

This is partially in line with studies proposing that conflict processing may rely on a hybrid architecture. Yet, what qualifies as hybrid differs considerably across accounts. For example, Li and colleagues (2017) suggest that cognitive control draws on both a domain-general system, supported by the frontoparietal and cingulo-opercular networks, and domain-specific systems that vary across tasks. However, such a proposal does not readily accommodate our findings. If each task recruited a distinct domain-specific module in addition to a shared module, one would expect successful cross-task decoding across all task combinations, which we did not observe.

A complementary hybrid perspective comes from the distributed control system account (Zink et al., 2021), which proposes that conflict processing emerges from flexible interactions within widely distributed networks rather than from fixed, task-invariant regions. This view aligns with evidence that the frontoparietal network operates as a flexible hub that reconfigures its connectivity to meet changing task demands (Cole et al., 2013; Menon & D’Esposito, 2022; Power & Petersen, 2013). Although EEG cannot directly assess connectivity, scalp-level decoding likely reflects coordinated activity across such distributed systems. In this context, the robust transfer of conflict information between the Stroop Naming task and Change Detection task and Stroop Naming task and Simon task, suggests some overlapping network dynamics. Conversely, the absence of decoding generalization between the Simon task and Change Detection task points to more distinct configurations of neural engagement. Overall, the pattern is consistent with the idea that conflict tasks rely on shared but non-identical distributed control networks.

A complementary hybrid perspective that aligns well with the observed cross-task decoding results is provided by the “cognitive space” framework. This account proposes that different forms of conflict occupy locations within a structured representational space defined by their similarity relations (e.g., Yang et al., 2024). Within this framework, conflict processing is not characterized by a strict domain-general versus domain-specific divide; rather, it emerges from graded representational distances among conflict types. Tasks that are located closer in this cognitive space share more similar underlying control demands, whereas more distant tasks rely on increasingly distinct processing configurations. A related argument has been advanced by Becker and colleagues (2024), who suggest that differences between conflict tasks can be captured along continuous dimensions of task complexity. More complex conflicts require more diverse or flexible control operations, leading to more heterogeneous patterns of action implementation. Together, these perspectives converge on a broader conceptualization in which the mechanisms underlying conflict processing vary along continuous dimensions, rather than falling into discrete categories.

Viewed through this lens, the present findings offer empirical support for treating domain-generality and domain-specificity as endpoints on a continuum of similarity and shared neural mechanisms. The asymmetries in cross-task decoding suggest that some tasks occupy relatively proximal positions in this continuum, whereas others lie farther apart and exhibit little or no generalization. Future studies using tools optimized for assessing large-scale neural dynamics across diverse conflict tasks will be essential for testing these hypotheses more directly. Such work will further refine the emerging picture of conflict processing as a graded phenomenon, rather than a dichotomy between domain-general and domain-specific mechanisms.

### Conclusions

To summarize, we investigated whether the neural mechanisms underlying conflict processing are domain-general, conflict-specific, or whether an account involving partially overlapping mechanisms better captures these neural dynamics. We analyzed EEG data from 507 participants who completed three conflict tasks: a Change Detection task, the Simon task, and the Stroop task. Our results showed that while conflict signals could be reliably tracked within each task, they only partially generalized across tasks, with shared conflict signals between the Stroop Naming task and Change Detection task, and Stroop Naming task and Simon task. These findings support the partially overlapping neural mechanisms account suggesting that these neural resources are neither entirely shared nor fully task specific.

## 4. Methods

### 4.1. Participants

The present study utilized data collected within the ongoing Dortmund Vital Study (Clinicaltrials.gov NCT05155397; see Gajewski et al., 2022). At the time the current analyses were conducted, data had been collected from 614 participants. The exclusion criteria of the Dortmund Vital Study included no history of significant medical conditions, including (i) neurological disorders (e.g., dementia, Parkinson’s disease, or stroke); (ii) cardiovascular diseases; (iii) bleeding disorders; (iv) cancer; (v) psychiatric conditions (e.g., schizophrenia, obsessive-compulsive disorder, anxiety disorders, or severe depression); (vi) eye conditions such as cataracts, glaucoma, or blindness. Additionally, participants with a history of head injuries, surgeries, implants and those with a reduced physical fitness or mobility were excluded. Finally, participants taking psychotropic drugs or neuroleptics were omitted from the study. However, individuals taking medications such as blood thinners, hormones, antihypertensives, or cholesterol-lowering drugs were eligible for inclusion. All participants had normal or corrected-to-normal vision and hearing.

Originally, data were collected from 614 participants, but 107 participants were excluded for various reasons; Sixty participants did not attend the second experimental session required to complete the Stroop task. Twenty-one participants had excessively noisy EEG recordings for at least one task, rendering their data unusable. Eleven participants failed the Ishihara color test, and three had vision impairments. One participant could not complete testing due to health issues, and five were excluded for not being able to complete the tasks. Additionally, two participants were excluded due to technical issues encountered during their experimental sessions. Finally, the data from four participants were excluded because they missed too many trials during the Simon task, rendering their EEG data unsuitable for the decoding analyses. The final sample included 507 participants (320 females, 187 males, age range: 20-70 years, *M_age_* = 43.58, *SD_age_* = 14.09).

All participants provided written informed consent prior to participation, and the study adhered to the ethical principles outlined in the Declaration of Helsinki. Ethical approval was obtained from the local ethics committee of the Leibniz Research Centre for Working Environment and Human Factors, Dortmund, Germany (approval number: A93). The four tasks included in the current study were completed in two different experimental sessions that spanned two days. Participants were compensated with 160 Euros for these two sessions.

### 4.2. Experimental procedure

Participants completed computer-based cognitive tasks, including a Change Detection task (Wascher & Beste, 2010), a Simon task (Hoppe et al., 2017; Simon, 1969), and a Stroop task (Stroop, 1935). The Change Detection task and Simon task were administered on the first day, whereas the Stroop task was conducted in a separate session on a different day. EEG signals were recorded throughout the completion of these tasks.

#### 4.2.1. Technical set-up – first session

Tasks were displayed on a 32-inch VSG monitor (Display++ LCD, M0250 & M0251) with a resolution of 1920×1080 pixels and a refresh rate of 100 Hz. Manual responses were captured using force-sensitive handles. The stimuli and presentation sequence were created using the FreePascal software. EEG data were collected using an Ag–AgCI active electrode EEG system (actiCap; Brain Products GmbH). The signal was sampled at a rate of 1000 Hz and filtered in real time with a 200 Hz low-pass filter. A 64-channel cap was used for data acquisition, with the FCz electrode serving as the reference electrode and Afz as the ground electrode.

#### 4.2.2. Technical set-up – second session

On the second day, participants performed the tasks on a 17-inch monitor (refresh rate: 100 Hz, resolution: 640 × 480 pixels) and were seated ∼70 cm away from the screen. For recording the EEG data, a 32-channel EEG system equipped with Ag–AgCl active electrodes (BioSemi B.V.) was utilized, with data sampled online at 2048 Hz. The BioSemi system incorporates a Common Mode Sense (CMS) active electrode and a Driven Right Leg (DRL) passive electrode, which together establish a feedback loop to regulate the subject’s average potential. The reference and ground electrodes are included within this CMS and DRL loop. Electrode placement followed the international 10–20 system, with impedances maintained below 10 kΩ during both experimental sessions.

#### 4.2.3. Change Detection task

Participants were shown a display (luminosity: 20cd/m^2^) with two bars (size: 1.35°×0.56° visual angle; color: CIE1931: 0.287, 0.312, 10-50 cd/m^2^, where the last parameter has been varied between 10-50). One of the bars was always on the left, while the other on the right of a central fixation dot (size: 0.3°×0.3°; distance between fixation dot and bar: 1.3° visual angle). After 50 milliseconds, a second display appeared introducing either a luminance change, an orientation change, or both. Luminance changes occurred either from 10 to 50 cd/m^2^ or from 50 to 10 cd/m^2^. Similarly, orientation changes could switch from horizontal to vertical or from vertical to horizontal (Wascher & Beste, 2010). Participants’ task was to indicate, via a left– or right-hand button press, whether the luminance change occurred on the left or right side. The luminance and orientation changes could appear either together or separately. Importantly, the task also included a condition in which only the orientation of the stimulus changed. However, these trials were omitted from the present analyses because they involve a qualitatively different form of conflict: orientation-only changes signal a “no-go”-type response requirement, where participants must withhold a response despite detecting a salient change. Including orientation-only trials as conflict trials would therefore conflate two distinct processes, response inhibition and stimulus–response conflict, which would complicate both interpretation and comparability with the Stroop task and Simon task. For this reason, we restricted analyses to trials where a luminance change required a response. Overall, trials were categorized as: (i) non-conflict trials, where only a luminance change occurred, or a luminance change occurred along with an orientation change in the same bar; (ii) conflict trials, in which the luminance change occurred in one bar while the orientation change occurred in the other bar.

#### 4.2.4. Simon task

Trials in the Simon task began with a central fixation cross (size: 0.3°×0.3°) and two placeholder dots (size: 0.15°×0.15°), one on each side of the fixation displayed for a variable interval between 500 and 800 milliseconds (background color: CIE1931: 0.287, 0.312, 10). Following this, participants were shown a stimulus display consisting of two shapes on either side of the fixation cross (distance between shape and central fixation: 2° visual angle). Participants were instructed to press a button when circles appeared and to withhold their response when a diamond was present. For the current study, we only consider trials, in which a response had to be made. Within the circles (diameter in visual angles: 1.1°), letters could be displayed, always appearing on one side of the fixation cross, while the other shape contained three horizontal lines (size of each line: 0.45°×0.07° visual angle) as a placeholder (color: CIE1931: 0.287, 0.312, 80). If participants saw the letter “H” (size: 0.536°×0.474° visual angle) they were supposed to respond with one hand (e.g., left), and if they saw the letter “N” (size: 0.495°×0.474° visual angle) they were required to respond with the other hand (e.g., right). Importantly, trials were categorized as: (i) non-conflict trials when the letter that instructed the participant to press the button with a particular hand (e.g., left) appeared on the corresponding side (e.g., left); (ii) conflict trials when the letter instructing the participant to press with a specific hand appeared on the opposite side (e.g., the letter for the left hand was on the right side).

#### 4.2.5. Stroop task

The task was divided into two main blocks. In the first block, participants’ task was to indicate the word’s meaning (i.e., read the color word), while in the second block, they had to report the color of the ink, in which the word is printed. The Reading block thus served as a baseline task, in which no conflict was involved, as participants only had to read the word. The trial started with a cue (square or diamond, size: 0.33°×3.03° visual angle) indicating whether it is a color reading or a color naming task. Following a 1000 ms inter-stimulus-interval, participants were displayed a color word (size: 0.57°×0.82° visual angle). The stimuli consisted of the German words “rot,” “grün,” “gelb,” “blau” for “red,” “green,” “yellow” and “blue” each displayed in one of these four colors. The color of the presented words was either compatible or incompatible with the word’s meaning. Half of the trials were compatible (e.g., the word “red” displayed with red ink–non-conflict trials), and the other half were incompatible (e.g., the word “red” in green color – conflict trials). To respond, participants used four buttons, each of which had an assigned color that was learnt before starting the session. For responses, the index and middle fingers of both hands were used. The color-button assignment was the same for all participants. Participants had 2500 milliseconds to respond. At the end of the trial, they received feedback: a plus sign for correct responses and a minus sign (size: 0.82°×0.82° visual angle) for incorrect responses. The response-cue interval was 1300 milliseconds and included the response feedback and a feedback delay. The instruction encouraged both quick and accurate responses.

### 4.3. Data analyses

#### 4.3.1. Behavioral analyses

As an initial step, we examined whether conflict processing was reflected behaviorally by calculating mean accuracy and response times for conflict and non-conflict trials across the four tasks. Data were first cleaned by removing trials with premature responses (i.e., button presses before the test screen) and excessively long response times. Specifically, we excluded trials with response times exceeding 1800 ms in the Change Detection task, 1500 ms in the Simon task, and 4000 ms in the Stroop task. We then compared accuracy and response times between conflict and non-conflict conditions using paired-sample t-tests. Additionally, for each task we computed conflict effects (non-conflict minus conflict accuracy and response times) and assessed their relationships across tasks using Spearman correlations.

#### 4.3.2. EEG preprocessing

Given the robustness of multivariate methods to noise (Carlson et al., 2020) minimal preprocessing was performed. Data were high-pass and low-pass filtered using a Hamming windowed sinc FIR filter and then downsampled to 250 Hz. For each task, epochs time-locked to the target stimulus were created: (i) Change Detection task: –500 to 2800 ms; (ii) Simon task: –500 to 2500 ms; (iii) both Stroop tasks: –400 to 2000 ms. At the end, baseline removal (0–200 ms) was applied. Preprocessing was conducted using functions from the EEGLAB toolbox (v2024.0; (Delorme & Makeig, 2004) implemented in MATLAB.

#### 4.3.3. Decoding analyses & statistics

To address the main research question, multivariate decoding analyses were performed using the CoSMoMVPA toolbox (Oosterhof et al., 2016) implemented in MATLAB. For all analyses, the classifier was trained and tested to distinguish between conflict and non-conflict trials. Therefore, the chance level was always set to 0.5 (i.e., 50%). Regularized linear discriminant analysis classifiers were employed. To ensure consistency and comparability of results, each dataset was standardized to include a common set of electrodes across all participants and sessions. Trials where participants failed to respond within the time limit were excluded. Table 3 summarizes the mean number of conflict and non-conflict trials in each task. Importantly, the apparent imbalance in the Change Detection task arises because we did not include the orientation-change trials in this analysis. Additionally, for all decoding analyses, we ensured equal numbers of trials per condition and across tasks, thereby eliminating imbalance as a potential confound.

**Table 3.** Average number of conflict and non-conflict trials across tasks.

| Task | Conflict | Non-conflict | *Conflict, only correct* | *Non-conflict, only correct* |
| --- | --- | --- | --- | --- |
| Change Detection | 75.96 trials | 162.99 trials | 53.01 trials | 145.29 trials |
| Simon | 226.37 trials | 228.55 trials | 209.75 trials | 217.57 trials |
| Stroop Reading | 51.90 trials | 51.90 trials | 50.60 trials | 50.77 trials |
| Stroop Naming | 51.69 trials | 51.80 trials | 48.14 trials | 50.69 trials |

##### 4.3.3.1. Within-task decoding – time domain analysis

As the first step in the analysis, we performed within-task decoding, where the classifier was both trained and tested using trials from the same task. We employed a 10-fold cross-validation approach, training the classifier on 90% of the data and testing it on the remaining 10%. This process was repeated across all folds, ensuring each data segment served as the test set once. To maintain class-balance, the `cosmo_balance_partitions` function was used to ensure equal representation of conflict and non-conflict trials in both the training and testing sets. The classifier was trained and tested independently at each time point, providing precise temporal resolution of decoding accuracy (Grootswagers et al., 2017). For each time point, decoding performance was calculated as the proportion of correctly classified trials, reflecting the classifier’s ability to differentiate between conflict and non-conflict conditions. We also performed a searchlight analysis. Here, we used electrode values as features, so the analysis was performed individually for each electrode and timepoint, yielding topographical maps at the following key timepoints: 252, 500, 752, 1000, 1500 ms. Statistical analyses were conducted using the `cosmo_montecarlo_cluster_stat`, which incorporates Threshold-Free Cluster Enhancement (Smith & Nichols, 2009) and Monte Carlo-based permutation testing (Maris & Oostenveld, 2007) to account for multiple comparisons (Oosterhof et al., 2016).

##### 4.3.3.2. Within-task decoding – frequency-domain analysis

In a second step, we implemented an alternative within-task decoding analysis, which uses power values from different frequencies as a feature alongside electrodes. We implemented this second within-task decoding procedure to determine whether power values from specific frequency ranges significantly contribute to decoding accuracy (Hanslmayr et al., 2008; Nigbur et al., 2011). This approach excludes the temporal dimension by applying a fast Fourier transform to the 0–1000 ms window following stimulus presentation, converting raw EEG signals into power at each frequency value. Power values within the 1–50 Hz range were extracted and used as features in the decoding analysis alongside electrode data, resulting in a single decoding accuracy value per participant. As in the previous decoding analysis, we employed a 10-fold cross-validation procedure, ensuring equal representation of conflict and non-conflict trials in both training and testing sets. Before running the decoding procedure, values underwent z-transformation. Finally, classifier performance was quantified as the proportion of correctly classified trials. For the searchlight analysis, we followed the same procedure, with one key difference: instead of using electrodes and frequency values as features, the analysis was conducted separately for each electrode and frequency combination. First, we plotted the average decoding accuracy across all electrodes for the frequency range 1-50 Hz. Additionally, topographies are also shown for a set of key frequencies: 3, 5, 7, 10, 15, 20, 25, 30 Hz.

To evaluate the statistical reliability of the results, we generated a null distribution for each task. Conflict and non-conflict labels were randomly permuted 100 times per participant, generating 100 decoding accuracy values per participant. From these, one decoding accuracy value per participant was randomly selected, and the mean decoding accuracy across participants was calculated (Stelzer et al., 2013). This process was repeated 10,000 times to construct the null distribution. Finally, we compared the observed mean decoding accuracy to this null distribution, determining the percentile at which it fell. A percentile value falling below the 95th percentile is interpreted as evidence for the null hypothesis, while values above 95 represent evidence for significant above-chance decoding.

##### 4.3.3.3. Cross-task decoding – temporal generalization analysis

The cross-task decoding procedure was designed to examine whether neural patterns associated with conflict and non-conflict trials could generalize across time and different cognitive tasks. Notably, the Stroop Reading task was excluded from this set of analyses due to its lack of robust evidence for within-task decoding. The cross-task decoding procedure was modeled after the steps used in the within-task decoding analysis applied to the time domain. Specifically, the aim was to examine whether conflict decoding generalizes across different timepoints of different tasks (King & Dehaene, 2014). Thus, the classifier was trained to discriminate between conflict vs. non-conflict trials at each time point of one task and then it was tested at all other time points of the other task.

### 4.4. Bayesian statistics

Bayesian analyses were carried out using the MATLAB toolbox for Bayes factor analysis (Krekelberg, 2024). Within the Bayesian framework, Bayes factors indicate the strength of evidence supporting either the alternative or null hypothesis. BF_10_ quantifies the evidence for an effect or condition difference, while BF_01_ (1/BF_10_) quantifies the evidence for the absence of such an effect. Bayes factors less than 3 are considered weak / inconclusive evidence, those between 3 and 20 as positive or substantial, between 20 and 150 as strong, and values above 150 as very strong evidence (Wagenmakers, 2007).

### Author contribution – CrediT statement

M.S.: Conceptualization, Data Curation, Software, Methodology, Validation, Formal Analysis, Visualization, Project Administration, Writing – Original Draft, Writing – Review & Editing; M.V.: Methodology, Writing – Review & Editing; E.W.: Data Curation, Project Administration, Funding acquisition, Writing – Review & Editing. P.D.G.: Data Curation, Project Administration, Writing – Review & Editing. T.G.: Conceptualization, Software, Methodology, Writing – Review & Editing. All authors have read and agreed to the published version of the manuscript.

### Open practices and data availability statement

A sample dataset and all the scripts used for the reported analyses are publicly available on Open Science Framework (OSF): https://doi.org/10.17605/OSF.IO/EDTNM. Access to the full dataset can be requested by contacting the Leibniz Research Centre for Working Environment and Human Factors, Dortmund, Germany according to the research data management rules outlined in the protocol of the Dortmund Vital Study (Gajewski et al., 2022).

## Acknowledgments

We wish to thank all participants in the study and our colleagues at IfADo for excellent technical support, and numerous student assistants for conducting telephone interviews, organizing and conducting the testing and data pre-processing. The study was endorsed by the German Center for Mental Health (DZPG). Additionally, we would like to thank Daniel Schneider, Stephan Getzmann, Emad Alyan, and Stefan Arnau for their valuable input and comments.

## References

1. Aczel, B., Kovacs, M., Bognar, M., Palfi, B., Hartanto, A., Onie, S., Tiong, L. E., & Evans, T. R. (2021). Is there evidence for cross-domain congruency sequence effect? A replication of Kan et al.(2013). Royal Society Open Science, 8(3), 191353.

2. Akçay, Ç., & Hazeltine, E. (2011). Domain-specific conflict adaptation without feature repetitions. Psychonomic Bulletin & Review, 18(3), 505–511. 10.3758/s13423-011-0084-y

3. Allefeld, C., Görgen, K., & Haynes, J.-D. (2016). Valid population inference for information-based imaging: From the second-level t –test to prevalence inference. NeuroImage, 141, 378–392. 10.1016/j.neuroimage.2016.07.040

4. Becker, D., Bijleveld, E., Braem, S., Fröber, K., Götz, F. J., Kleiman, T., Körner, A., Pfister, R., Reiter, A. M. F., Saunders, B., Schneider, I. K., Soutschek, A., Van Steenbergen, H., & Dignath, D. (2024). An integrative framework of conflict and control. Trends in Cognitive Sciences, 28(8), 757–768. 10.1016/j.tics.2024.07.002

5. Boy, F., Husain, M., & Sumner, P. (2010). Unconscious inhibition separates two forms of cognitive control. Proceedings of the National Academy of Sciences, 107(24), 11134–11139. 10.1073/pnas.1001925107

6. Braem, S., Abrahamse, E. L., Duthoo, W., & Notebaert, W. (2014). What determines the specificity of conflict adaptation? A review, critical analysis, and proposed synthesis. Frontiers in Psychology, 5. 10.3389/fpsyg.2014.01134

7. Carlson, T. A., Grootswagers, T., & Robinson, A. K. (2020). An Introduction to Time-Resolved Decoding Analysis for M/EEG. In D. Poeppel, G. R. Mangun, & M. S. Gazzaniga (Eds.), The Cognitive Neurosciences (p. 0). The MIT Press. 10.7551/mitpress/11442.003.0075

8. Cespón, J., Hommel, B., Korsch, M., & Galashan, D. (2020). The neurocognitive underpinnings of the Simon effect: An integrative review of current research. *Cognitive, Affective*, & Behavioral Neuroscience, 20(6), 1133–1172. 10.3758/s13415-020-00836-y

9. Chen, T., Becker, B., Camilleri, J., Wang, L., Yu, S., Eickhoff, S. B., & Feng, C. (2018). A domain-general brain network underlying emotional and cognitive interference processing: Evidence from coordinate-based and functional connectivity meta-analyses. Brain Structure and Function, 223(8), 3813–3840. 10.1007/s00429-018-1727-9

10. Cole, M. W., Reynolds, J. R., Power, J. D., Repovs, G., Anticevic, A., & Braver, T. S. (2013). Multi-task connectivity reveals flexible hubs for adaptive task control. Nature Neuroscience, 16(9), 1348–1355. 10.1038/nn.3470

11. Delorme, A., & Makeig, S. (2004). EEGLAB: an open source toolbox for analysis of single-trial EEG dynamics including independent component analysis. Journal of Neuroscience Methods, 134(1), 9–21. 10.1016/j.jneumeth.2003.10.009

12. Dudschig, C. (2022). Are control processes domain-general? A replication of ‘To adapt or not to adapt? The question of domain-general cognitive control’ (Kan et al. 2013). Royal Society Open Science, 9(7). 10.1098/rsos.210550

13. Egner, T. (2008). Multiple conflict-driven control mechanisms in the human brain. Trends in Cognitive Sciences, 12(10), 374–380. 10.1016/j.tics.2008.07.001

14. Egner, T., Delano, M., & Hirsch, J. (2007). Separate conflict-specific cognitive control mechanisms in the human brain. NeuroImage, 35(2), 940–948. 10.1016/j.neuroimage.2006.11.061

15. Fernandez-Duque, D., & Knight, M. (2008). Cognitive control: Dynamic, sustained, and voluntary influences. Journal of Experimental Psychology: Human Perception and Performance, 34(2), 340–355. 10.1037/0096-1523.34.2.340

16. Forster, S. E., & Cho, R. Y. (2014). Context Specificity of Post-Error and Post-Conflict Cognitive Control Adjustments. PLoS ONE, 9(3), e90281. 10.1371/journal.pone.0090281

17. Freitas, A. L., Bahar, M., Yang, S., & Banai, R. (2007). Contextual Adjustments in Cognitive Control Across Tasks. Psychological Science, 18(12), 1040–1043. 10.1111/j.1467-9280.2007.02022.x

18. Freitas, A. L., & Clark, S. L. (2015). Generality and specificity in cognitive control: Conflict adaptation within and across selective-attention tasks but not across selective-attention and Simon tasks. Psychological Research, 79(1), 143–162. 10.1007/s00426-014-0540-1

19. Fu, Z., Beam, D., Chung, J. M., Reed, C. M., Mamelak, A. N., Adolphs, R., & Rutishauser, U. (2022). The geometry of domain-general performance monitoring in the human medial frontal cortex. Science, 376(6593). 10.1126/science.abm9922

20. Funes, M. J., Lupiáñez, J., & Humphreys, G. (2010). Analyzing the generality of conflict adaptation effects. Journal of Experimental Psychology: Human Perception and Performance, 36(1), 147–161. 10.1037/a0017598

21. Gajewski, P. D., & Falkenstein, M. (2015). Long-term habitual physical activity is associated with lower distractibility in a Stroop interference task in aging: Behavioral and ERP evidence. Brain and Cognition, 98, 87–101. 10.1016/j.bandc.2015.06.004

22. Gajewski, P. D., Falkenstein, M., Thönes, S., & Wascher, E. (2020). Stroop task performance across the lifespan: High cognitive reserve in older age is associated with enhanced proactive and reactive interference control. NeuroImage, 207, 116430. 10.1016/j.neuroimage.2019.116430

23. Gajewski, P. D., Getzmann, S., Bröde, P., Burke, M., Cadenas, C., Capellino, S., Claus, M., Genç, E., Golka, K., Hengstler, J. G., Kleinsorge, T., Marchan, R., Nitsche, M. A., Reinders, J., Van Thriel, C., Watzl, C., & Wascher, E. (2022). Impact of Biological and Lifestyle Factors on Cognitive Aging and Work Ability in the Dortmund Vital Study: Protocol of an Interdisciplinary, Cross-sectional, and Longitudinal Study. JMIR Research Protocols, 11(3), e32352. 10.2196/32352

24. Grootswagers, T., Wardle, S. G., & Carlson, T. A. (2017). Decoding Dynamic Brain Patterns from Evoked Responses: A Tutorial on Multivariate Pattern Analysis Applied to Time Series Neuroimaging Data. Journal of Cognitive Neuroscience, 29(4), 677–697. 10.1162/jocn_a_01068

25. Hanslmayr, S., Pastötter, B., Bäuml, K.-H., Gruber, S., Wimber, M., & Klimesch, W. (2008). The Electrophysiological Dynamics of Interference during the Stroop Task. Journal of Cognitive Neuroscience, 20(2), 215–225. 10.1162/jocn.2008.20020

26. Hebart, M. N., & Baker, C. I. (2018). Deconstructing multivariate decoding for the study of brain function. NeuroImage, 180, 4–18. 10.1016/j.neuroimage.2017.08.005

27. Heidlmayr, K., Kihlstedt, M., & Isel, F. (2020). A review on the electroencephalography markers of Stroop executive control processes. Brain and Cognition, 146, 105637. 10.1016/j.bandc.2020.105637

28. Hommel, B. (2011). The Simon effect as tool and heuristic. Acta Psychologica, 136(2), 189–202. 10.1016/j.actpsy.2010.04.011

29. Hoppe, K., Küper, K., & Wascher, E. (2017). Sequential Modulations in a Combined Horizontal and Vertical Simon Task: Is There ERP Evidence for Feature Integration Effects? Frontiers in Psychology, 8, 1094. 10.3389/fpsyg.2017.01094

30. Ila, A. B., & Polich, J. (1999). P300 and response time from a manual Stroop task. Clinical Neurophysiology, 110(2), 367–373. 10.1016/S0168-5597(98)00053-7

31. Jiang, J., & Egner, T. (2014). Using Neural Pattern Classifiers to Quantify the Modularity of Conflict-Control Mechanisms in the Human Brain. Cerebral Cortex, 24(7), 1793–1805. 10.1093/cercor/bht029

32. Kan, I. P., Teubner-Rhodes, S., Drummey, A. B., Nutile, L., Krupa, L., & Novick, J. M. (2013). To adapt or not to adapt: The question of domain-general cognitive control. Cognition, 129(3), 637–651. 10.1016/j.cognition.2013.09.001

33. Kim, C., Chung, C., & Kim, J. (2012). Conflict adjustment through domain-specific multiple cognitive control mechanisms. Brain Research, 1444, 55–64. 10.1016/j.brainres.2012.01.023

34. King, J.-R., & Dehaene, S. (2014). Characterizing the dynamics of mental representations: The temporal generalization method. Trends in Cognitive Sciences, 18(4), 203–210. 10.1016/j.tics.2014.01.002

35. Kleiman, T., Hassin, R. R., & Trope, Y. (2014). The control-freak mind: Stereotypical biases are eliminated following conflict-activated cognitive control. Journal of Experimental Psychology: General, 143(2), 498–503. 10.1037/a0033047

36. Krekelberg, B. (2024). *Matlab Toolbox for Bayes Factor Analysis* (Version v3.0) [Computer software]. Zenodo. 10.5281/ZENODO.13744717

37. Kunde, W., Augst, S., & Kleinsorge, T. (2012). Adaptation to (non)valent task disturbance. *Cognitive, Affective*, & Behavioral Neuroscience, 12(4), 644–660. 10.3758/s13415-012-0116-8

38. Kunde, W., & Stöcker, C. (2002). A Simon effect for stimulus-response duration. The Quarterly Journal of Experimental Psychology Section A, 55(2), 581–592. 10.1080/02724980143000433

39. Leuthold, H. (2011). The Simon effect in cognitive electrophysiology: A short review. Acta Psychologica, 136(2), 203–211. 10.1016/j.actpsy.2010.08.001

40. Li, H., Liu, N., Li, Y., Weidner, R., Fink, G. R., & Chen, Q. (2019). The Simon Effect Based on Allocentric and Egocentric Reference Frame: Common and Specific Neural Correlates. Scientific Reports, 9(1), 13727. 10.1038/s41598-019-49990-5

41. Li, Q., Yang, G., Li, Z., Qi, Y., Cole, M. W., & Liu, X. (2017). Conflict detection and resolution rely on a combination of common and distinct cognitive control networks. Neuroscience & Biobehavioral Reviews, 83, 123–131. 10.1016/j.neubiorev.2017.09.032

42. MacLeod, C. M. (1991). Half a century of research on the Stroop effect: An integrative review. Psychological Bulletin, 109(2), 163.

43. Maris, E., & Oostenveld, R. (2007). Nonparametric statistical testing of EEG– and MEG-data. Journal of Neuroscience Methods, 164(1), 177–190. 10.1016/j.jneumeth.2007.03.024

44. Menon, V., & D’Esposito, M. (2022). The role of PFC networks in cognitive control and executive function. Neuropsychopharmacology, 47(1), 90–103. 10.1038/s41386-021-01152-w

45. Nigbur, R., Ivanova, G., & Stürmer, B. (2011). Theta power as a marker for cognitive interference. Clinical Neurophysiology, 122(11), 2185–2194. 10.1016/j.clinph.2011.03.030

46. Oosterhof, N. N., Connolly, A. C., & Haxby, J. V. (2016). CoSMoMVPA: Multi-Modal Multivariate Pattern Analysis of Neuroimaging Data in Matlab/GNU Octave. Frontiers in Neuroinformatics, 10. https://www.frontiersin.org/journals/neuroinformatics/articles/10.3389/fninf.2016.00027

47. Parris, B. A., Hasshim, N., Wadsley, M., Augustinova, M., & Ferrand, L. (2022). The loci of Stroop effects: A critical review of methods and evidence for levels of processing contributing to color-word Stroop effects and the implications for the loci of attentional selection. Psychological Research, 86(4), 1029–1053. 10.1007/s00426-021-01554-x

48. Parris, B. A., Wadsley, M. G., Hasshim, N., Benattayallah, A., Augustinova, M., & Ferrand, L. (2019). An fMRI Study of Response and Semantic Conflict in the Stroop Task. Frontiers in Psychology, 10, 2426. 10.3389/fpsyg.2019.02426

49. Polich, J. (2007). Updating P300: An integrative theory of P3a and P3b. Clinical Neurophysiology, 118(10), 2128–2148. 10.1016/j.clinph.2007.04.019

50. Power, J. D., & Petersen, S. E. (2013). Control-related systems in the human brain. Current Opinion in Neurobiology, 23(2), 223–228. 10.1016/j.conb.2012.12.009

51. Rünger, D., Schwager, S., & Frensch, P. A. (2010). Across-task conflict regulation: A replication failure. Journal of Experimental Psychology: Human Perception and Performance, 36(1), 136–146. 10.1037/a0017172

52. Schlaghecken, F., Refaat, M., & Maylor, E. A. (2011). Multiple systems for cognitive control: Evidence from a hybrid prime-Simon task. Journal of Experimental Psychology: Human Perception and Performance, 37(5), 1542–1553. 10.1037/a0024327

53. Schmidt, J. R., & Weissman, D. H. (2014). Congruency Sequence Effects without Feature Integration or Contingency Learning Confounds. PLoS ONE, 9(7), e102337. 10.1371/journal.pone.0102337

54. Schneider, D., Beste, C., & Wascher, E. (2012a). Attentional Capture by Irrelevant Transients Leads to Perceptual Errors in a Competitive Change Detection Task. Frontiers in Psychology, 3. 10.3389/fpsyg.2012.00164

55. Schneider, D., Beste, C., & Wascher, E. (2012b). On the time course of bottom-up and top-down processes in selective visual attention: An EEG study. Psychophysiology, 49(11), 1660–1671. 10.1111/j.1469-8986.2012.01462.x

56. Schneider, D., Hoffmann, S., & Wascher, E. (2014). Sustained posterior contralateral activity indicates re-entrant target processing in visual change detection: An EEG study. Frontiers in Human Neuroscience, 8. 10.3389/fnhum.2014.00247

57. Schneider, D., & Wascher, E. (2013). Mechanisms of target localization in visual change detection: An interplay of gating and filtering. Behavioural Brain Research, 256, 311–319. 10.1016/j.bbr.2013.08.046

58. Simon, J. R. (1969). Reactions toward the source of stimulation. Journal of Experimental Psychology, 81(1), 174–176. 10.1037/h0027448

59. Smith, S., & Nichols, T. (2009). Threshold-free cluster enhancement: Addressing problems of smoothing, threshold dependence and localisation in cluster inference. NeuroImage, 44(1), 83–98. 10.1016/j.neuroimage.2008.03.061

60. Stelzer, J., Chen, Y., & Turner, R. (2013). Statistical inference and multiple testing correction in classification-based multi-voxel pattern analysis (MVPA): Random permutations and cluster size control. NeuroImage, 65, 69–82. 10.1016/j.neuroimage.2012.09.063

61. Stroop, J. R. (1935). Studies of interference in serial verbal reactions. Journal of Experimental Psychology, 18(6), 643–662. 10.1037/h0054651

62. Ullsperger, M., & Von Cramon, D. Y. (2006). The Role of Intact Frontostriatal Circuits in Error Processing. Journal of Cognitive Neuroscience, 18(4), 651–664. 10.1162/jocn.2006.18.4.651

63. Vanveen, V., & Carter, C. (2002). The anterior cingulate as a conflict monitor: fMRI and ERP studies. Physiology & Behavior, 77(4–5), 477–482. 10.1016/S0031-9384(02)00930-7

64. Verbruggen, F., Liefooghe, B., Notebaert, W., & Vandierendonck, A. (2005). Effects of stimulus–stimulus compatibility and stimulus–response compatibility on response inhibition. Acta Psychologica, 120(3), 307–326. 10.1016/j.actpsy.2005.05.003

65. Vermeylen, L., Wisniewski, D., González-García, C., Hoofs, V., Notebaert, W., & Braem, S. (2020). Shared Neural Representations of Cognitive Conflict and Negative Affect in the Medial Frontal Cortex. The Journal of Neuroscience, 40(45), 8715–8725. 10.1523/jneurosci.1744-20.2020

66. Wagenmakers, E.-J. (2007). A practical solution to the pervasive problems ofp values. Psychonomic Bulletin & Review, 14(5), 779–804. 10.3758/BF03194105

67. Wascher, E., & Beste, C. (2010). Tuning Perceptual Competition. Journal of Neurophysiology, 103(2), 1057–1065. 10.1152/jn.00376.2009

68. Wendt, M., Kluwe, R. H., & Peters, A. (2006). Sequential modulations of interference evoked by processing task-irrelevant stimulus features. Journal of Experimental Psychology: Human Perception and Performance, 32(3), 644–667. 10.1037/0096-1523.32.3.644

69. Wühr, P., Duthoo, W., & Notebaert, W. (2015). Generalizing attentional control across dimensions and tasks: Evidence from transfer of proportion-congruent effects. Quarterly Journal of Experimental Psychology, 68(4), 779–801. 10.1080/17470218.2014.966729

70. Yang, G., Wu, H., Li, Q., Liu, X., Fu, Z., & Jiang, J. (2024). Dorsolateral prefrontal activity supports a cognitive space organization of cognitive control. eLife, 12. 10.7554/elife.87126

71. Yeung, N., & Cohen, J. D. (2006). The Impact of Cognitive Deficits on Conflict Monitoring: Predictable Dissociations Between the Error-Related Negativity and N2. Psychological Science, 17(2), 164–171. 10.1111/j.1467-9280.2006.01680.x

72. Zhu, D., Wang, X., Zhao, E., Nozari, N., Notebaert, W., & Braem, S. (2025). Cognitive control is task specific: Further evidence against the idea of domain-general conflict adaptation. *Journal of Experimental Psychology: Learning*, Memory, and Cognition.

73. Zink, N., Lenartowicz, A., & Markett, S. (2021). A new era for executive function research: On the transition from centralized to distributed executive functioning. Neuroscience & Biobehavioral Reviews, 124, 235–244. 10.1016/j.neubiorev.2021.02.011

74. Zmigrod, S., Zmigrod, L., & Hommel, B. (2016). Transcranial direct current stimulation (tDCS) over the right dorsolateral prefrontal cortex affects stimulus conflict but not response conflict. Neuroscience, 322, 320–325. 10.1016/j.neuroscience.2016.02.046

