## Supplementary materials for "Multiple Partially Overlapping Neural Modules Orchestrate Conflict Processing"

1. **Successful cross-temporal cross-task decoding of response hand**

As a control analysis, we tested whether the lack of cross-task conflict decoding summarized in section 2.4 can attributed to confounding factors that prevent the decoding of any feature. Using the same decoding procedure described in Section 2.4, we decoded response hand instead of conflict. As shown in Figure S1, response-hand information generalized robustly across all task combinations. When training on the Change Detection task and testing on the Simon task, we observed a significant cross-temporal generalization window from 184 to 1992 ms, which transferred to the -72-1992 ms interval in the Simon task. The reversed analysis (training on the Simon task, testing on the Change Detection task) yielded a significant window from -72-1992 ms that generalized to 200-1992 ms in the Change Detection task. Similarly, training on the Change Detection task and testing on the Stroop Naming task produced a generalizable signal emerging at 184 ms, lasting until 1992 ms, and transferring to -24-1992 ms in the Stroop Naming task. The reverse direction (training on the Stroop Naming task, testing on the Change Detection task) also revealed a robust generalization period from -8-1992 to 160-1992 ms. Finally, training on the Simon task yielded a significant cluster between 200-1992 ms that generalized to 240-1992 ms in the Stroop Naming task. The reversed analysis confirmed this pattern, with significant clusters observed at 200-1992 ms and 192-1992 ms.

2. **Cross-task decoding – evidence from the frequency domain**

Second, we conducted a cross-task decoding procedure, where both channel and frequency information between 1–50 Hz serving as input features (comparable to the within-task decoding procedure described in section 2.2). As shown in Figure S2, the results of this analysis revealed that the mean decoding accuracy values did not deviate from chance level, as indicated by the percentiles at which the mean decoding accuracy values are situated relative to the null distribution: (a) 4.99th percentile for the Change Detection task-Simon task; (b) 84.15th percentile for the Change Detection task-Stroop Naming task; (c) 4.90th percentile for the Simon task-Change Detection task; (d) 59.92th percentile for the Simon task-Stroop Naming task; (e) 14.72th percentile for the Stroop Naming task-Change Detection task; (f) 56.29th percentile for the Stroop Naming task-Simon task. Overall, these results suggest evidence for a lack of conflict signal generalizability across tasks.

Building on the within-task results, which suggested that decoding accuracy might be driven by theta-band power (3–7 Hz; cf., Hanslmayr et al., 2008; Nigbur et al., 2011), we refined the analysis by restricting input features to this frequency band. However, as illustrated in Figure

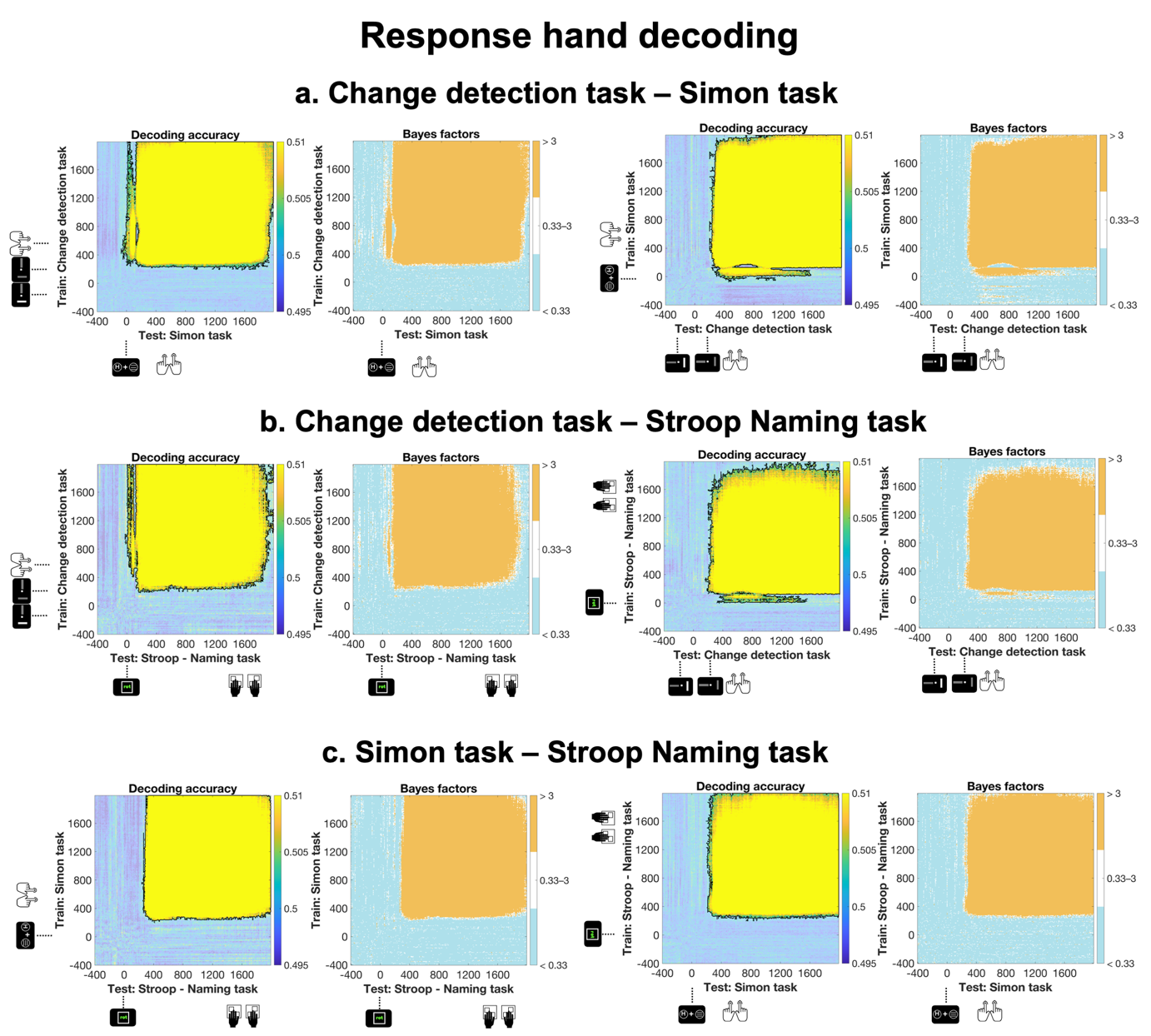

**Figure S1. Temporal generalization of response hand decoding across tasks.** Mean temporal generalization matrices showing cross-task decoding accuracy (left panels) and corresponding Bayes Factors (right panels) for all task pairings: (a) Change Detection task and Simon task, (b) Change Detection task and Stroop Naming task, and (c) Simon task and Stroop Naming task. In all decoding accuracy plots, light blue indicates chance-level temporal generalization accuracy, whereas warmer colors (yellow to orange) reflect above-chance decoding performance. Significant clusters, identified via cluster-based permutation testing, are outlined by black contours in the decoding accuracy plots. In all Bayes Factor plots, orange indicates Bayes Factors above 3 (at least substantial evidence for the alternative hypothesis), white depicts inconclusive evidence (Bayes factors between 0.33-3) and cooler blue shades indicate Bayes Factors below 3 (at least substantial evidence for the null hypothesis).

4b, this adjustment produced the same pattern of results: evidence was found for chance-level decoding performance across all comparisons: (a) 43.54th percentile for the Change Detection task-Simon task; (b) 49.26th percentile for the Change Detection task-Stroop Naming task; (c) 16.08th percentile for the Simon task-Change Detection task; (d) 10.62th percentile for the Simon task-Stroop Naming task; (e) 85.23th percentile for the Stroop Naming task-Change Detection task; (f) 77.67th percentile for the Stroop Naming task-Simon task.

To further test the robustness of these findings, we employed a third approach: a leave-one-task-out decoding procedure. In this method, a linear model was trained on all tasks except one, which was then used as the testing set. This approach was chosen to construct a more robust

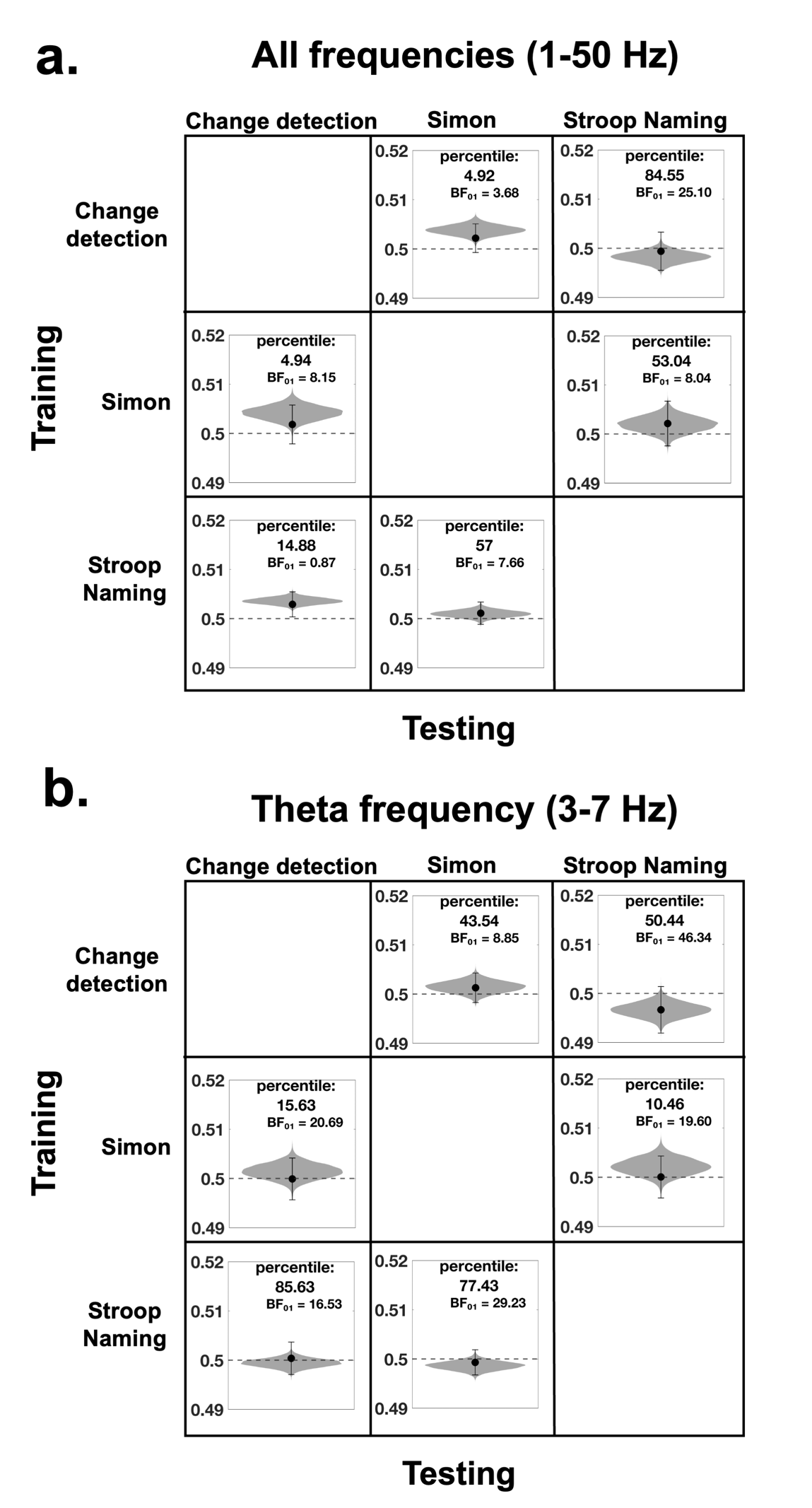

**Figure S2. Cross-task decoding results.** Both panels display the comparison of mean decoding accuracy relative to the null distribution across different training-testing pairs. Additionally, each sub-figure contains the percentile, where the mean decoding accuracy falls relative to the null distribution. Panel (a) represents results across all frequency ranges, and panel (b) focuses on the theta frequency range (3–7 Hz). The dashed gray line represents chance level, and the violin plot illustrates the null distribution generated by permuting conflict labels within each participant, with 10,000 permutations randomly sampled and averaged. As explained in Section 2.2, the null distribution values are not symmetrically centered around the chance level. The dot represents the mean decoding accuracy, and the error bar indicates the 95% confidence intervals of the mean.

model, potentially better suited to detect a task-general conflict processing signal. The statistical procedure was identical to that used in the within-task analyses described in Section 2.2. Specifically, we constructed a null distribution and identified the percentile at which the average decoding accuracy was located. If this percentile fell below the 95th, we interpreted it as support for the null hypothesis. Conversely, percentiles above the 95th were taken as evidence of significantly above-chance decoding performance. The results corroborated the earlier findings: cross-task decoding accuracy remained at chance level, providing strong evidence for the null hypothesis. Specifically, the average decoding accuracy was positioned at the 86.97th percentile of the null distribution.
